# Polymer biodegradation by *Halanaerobium* promotes reservoir souring during hydraulic fracturing

**DOI:** 10.1101/2023.06.16.545336

**Authors:** Gabrielle Scheffer, Anirban Chakraborty, Kaela K. Amundson, Rohan Khan, Michael J. Wilkins, Kelly Wrighton, Paul Evans, Casey R. J. Hubert

## Abstract

Hydraulically fractured shale reservoirs have facilitated studies of unexplored niches in the continental deep biosphere. Members of the genus *Halanaerobium* are ubiquitous in these environments. Polymers like guar gum used as gelling agents in hydraulic fracturing fluids are known to be fermentable substrates, but metabolic pathways encoding these processes have not been characterized. To explore this, produced water samples from the Permian Basin were incubated at 30°C to simulate above-ground storage conditions, and at 60°C to simulate subsurface reservoir conditions. Guar metabolism coincided with *Halanaerobium* enrichment only at 30°C, revealing genes for polymer biodegradation through the mixed-acid fermentation pathway in different metagenome-assembled genomes (MAGs). Whereas thiosulfate reduction to sulfide is often invoked to explain the dominance of *Halanaerobium* in these settings, *Halanaerobium* genomes did not uncover genes for this metabolism. Sulfide production was observed in 60°C incubations, with corresponding enrichment of *Desulfohalobium* and *Desulfovibrionaceae* that possess complete pathways for coupling mannose and acetate oxidation to sulfate reduction. These findings outline how production of fermentation intermediates (mannose, acetate) by *Halanaerobium* in topside settings can result in reservoir souring when these metabolites are introduced into the subsurface through produced water re-use.

**Importance:** Hydraulically fractured shale oil reservoirs are ideal for studying extremophiles such as thermohalophiles. During hydraulic fracturing, reservoir production water is stored in surface ponds prior to re-use. Microorganisms in these systems therefore need to withstand various environmental changes such as the swing between warm downhole oil reservoir temperatures and cooler surface conditions. This study follows this water cycle during fracking and the associated microbial metabolic potential. Of particular interest are members of the genus *Halanaerobium*, that have been reported to reduce thiosulfate contributing to souring of oil reservoirs. Here, we show that some *Halanaerobium* strains were unable to grow under oil reservoir temperatures and do not possess genes for thiosulfate reduction. Rather, it is likely that these organisms metabolize complex organics in fracking fluids at lower temperatures, thereby generating substrates that support reservoir souring by thermophilic sulfate-reducing bacteria at higher temperatures.

## Introduction

Studies of the terrestrial deep biosphere have advanced in recent years due to hydraulic fracturing for oil and gas production (1–3) showcasing polyextremophiles able to contend with high temperatures and salinities (4–9). Shale formations ranging from 45ºC to 190ºC (10) with salinities of 40,000 to 280,000 mg/L (11) are further impacted by organic polymers that are used in the hydraulic fracturing process. Common organic polymers used as gelling agents such as guar gum, carboxymethyl cellulose or polyacrylamide-based polymers have all been shown to be biodegradable as fermentable substrates or sources of nitrogen (12–15).

Among the most commonly detected microorganisms in hydraulically fractured shale reservoirs are members of the genus *Halanaerobium*. These microorganisms appear to be ubiquitous throughout most North American shale reservoir operations including the Marcellus, Barnett, Antrim and Haynesville shale systems (2). *Halanaerobium* spp. are known to ferment a wide range of carbohydrates, including guar gum (5, 16), but can also reduce elemental sulfur or thiosulfate to generate sulfide (6). Formation of precursors to sulfide biogenesis (e.g., zero-valent polysulfide or thiosulfate intermediates) depends on temperature, oxygen, pH, metals and the presence of other sulfur species (17). While some studies have measured very low to non-detectable levels of thiosulfate in shale reservoir systems (5, 6, 18, 19), others have amended sulfur species as electron acceptors at artificially high levels to provoke sulfide generation in enrichment cultures (20, 21). While the most well-understood paradigm for oil reservoir souring involves sulfate reduction, knowledge gaps persist regarding the mechanisms of souring in shale systems, where it is popular to invoke reduction of thiosulfate or elemental sulfur by *Halanaerobium in situ* (17, 20, 21). These proposals are supported by the physiology in pure cultures of *Halanaerobium* spp., including the type strain *H. congolense* (22).

Limited water sources in the vicinity of hydraulic fracturing operations lead many companies to reuse produced water from subsurface reservoirs to generate subsequent batches of fracturing fluids (23). As part of this process, microbial populations are exposed to different temperatures for prolonged periods between produced water stored in what is termed storage ponds at the surface and reintroduction into the reservoir. In the Permian Basin, shale formations tend to be around 60ºC, while surface storage ponds are commonly around 30ºC (24, 25). Little is known about the persistence and activity of microorganisms in topsides storage ponds, the populations that can survive for prolonged periods, the type of metabolic ability they have and whether that metabolism influences operations. The objectives of this study were to better understand how the genomic and biogeochemical dynamics between produced water storage ponds and hydraulically fractured oil reservoir conditions can contribute to carbon and sulfur cycling.

## Results

### Metabolic processes at 30°C and 60°C

In produced water enrichments at 60°C amended with glucose and a mix of six volatile fatty acids (VFA; acetate, butyrate, formate, lactate, propionate, succinate), the concentration of formate decreased between 14 and 21 days from 4.9 ± 0.1 to 3.6 ± 0.3 mM. Conversely, formate production was not observed in 30°C enrichments amended with guar gum (Figure 1A), where acetate levels increased to 2.8 ± 0.4 mM. A similar increase in acetate (up to 2.83 mM) was observed at 60°C with glucose and VFAs (Figure 1D). Acetate was also present in the incubations that did not receive VFA amendment (Fig. 1A, C) owing to being present at 2.1 mM in the original produced water (Table S1). Carbon dioxide production was the most pronounced within the enrichments at 30°C with guar gum, reaching a final concentration of 1.2 ± 0.1 mM (Figure 1A). Only the incubations at 60°C with glucose and VFA showed a clear decrease in sulfate concentration, which dropped from 8.7 ± 0.9 to 5.4 ± 0.4 mM over the 42-day incubation period, coinciding with an increase in sulfide from 1.4 ± 0.1 mM to 3.4 ± 0.5 mM (Figure 1D). Sulfate concentrations did not change significantly in the 60°C incubation amended with guar, or in either of the 30°C incubations (Figure 1A-C). No thiosulfate was detected within the microbial enrichments or the initial produced water through the experiments (Table S1). No metabolic activity was observed in any of the sterile or substrate-free controls.

**Figure 1:**
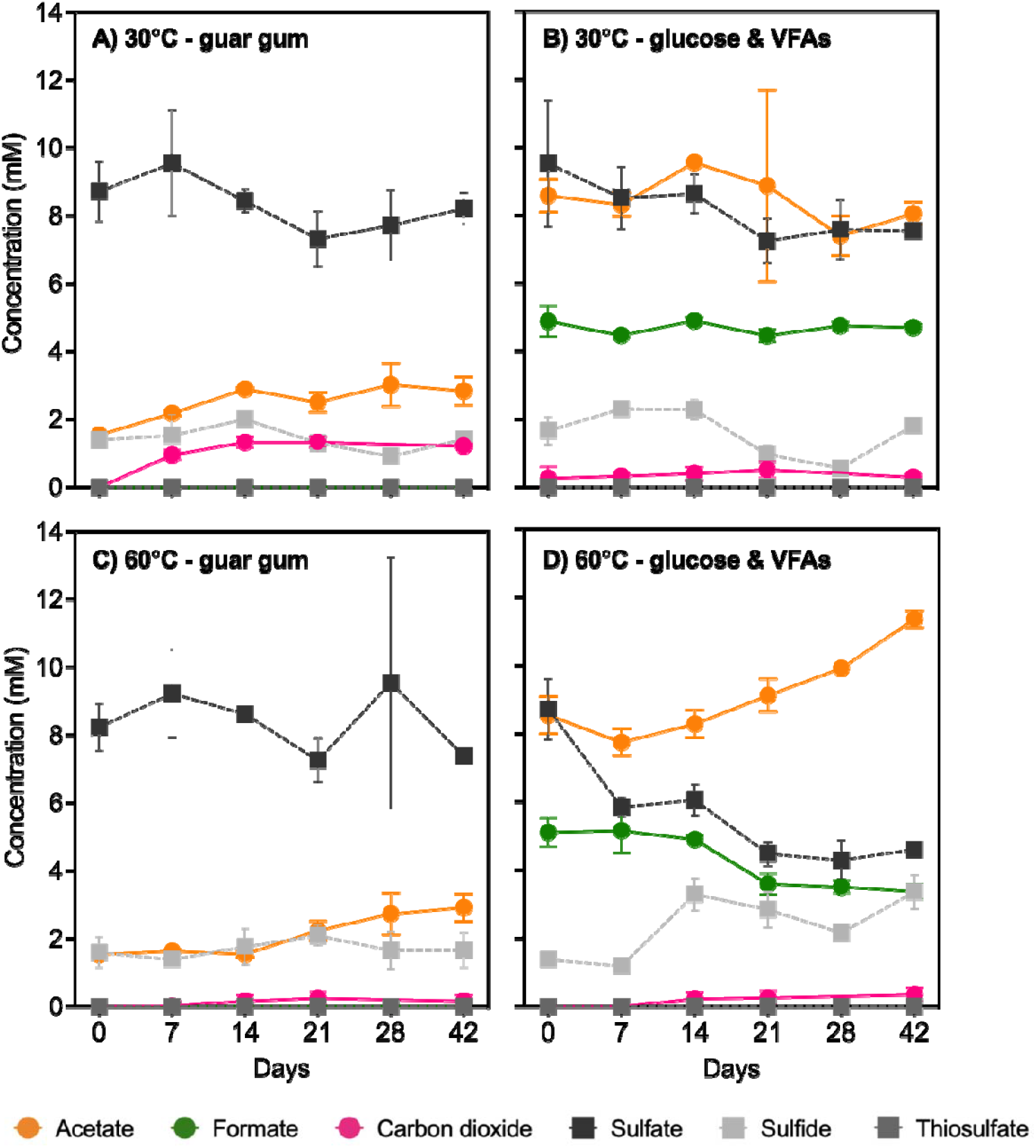
Concentrations of acetate, formate, carbon dioxide, sulfate and sulfide throughout 42-day incubations of produced water at 30°C and 60°C and amended with different substrates. Error bars represent standard deviation based on triplicate incubations for each condition. Carbon compounds are represented by dots connected with solid lines, and sulfur compounds are represented by squares connected with dashed lines.

### Microbial community composition

Microbial populations enriched during incubations at 30°C featured a higher relative abundance of *Halanaerobium* ASVs whereas at 60°C this group did not increase relative to initial levels detected in the produced water samples (Figure 2). Guar gum amendment at 30°C resulted in the highest relative abundance of *Halanaerobium*, with ASV3 reaching 47% after 42 days of incubation (Figure 2A). In addition, *Halanaerobium* ASV7 reached 10% after 21 days in the same 30°C incubation (Figure 2A), compared to other incubation conditions where it was detected at relative abundances of 1 to 7% compared to being 8% at day 0 (Figure 2B-D).

**Figure 2:**
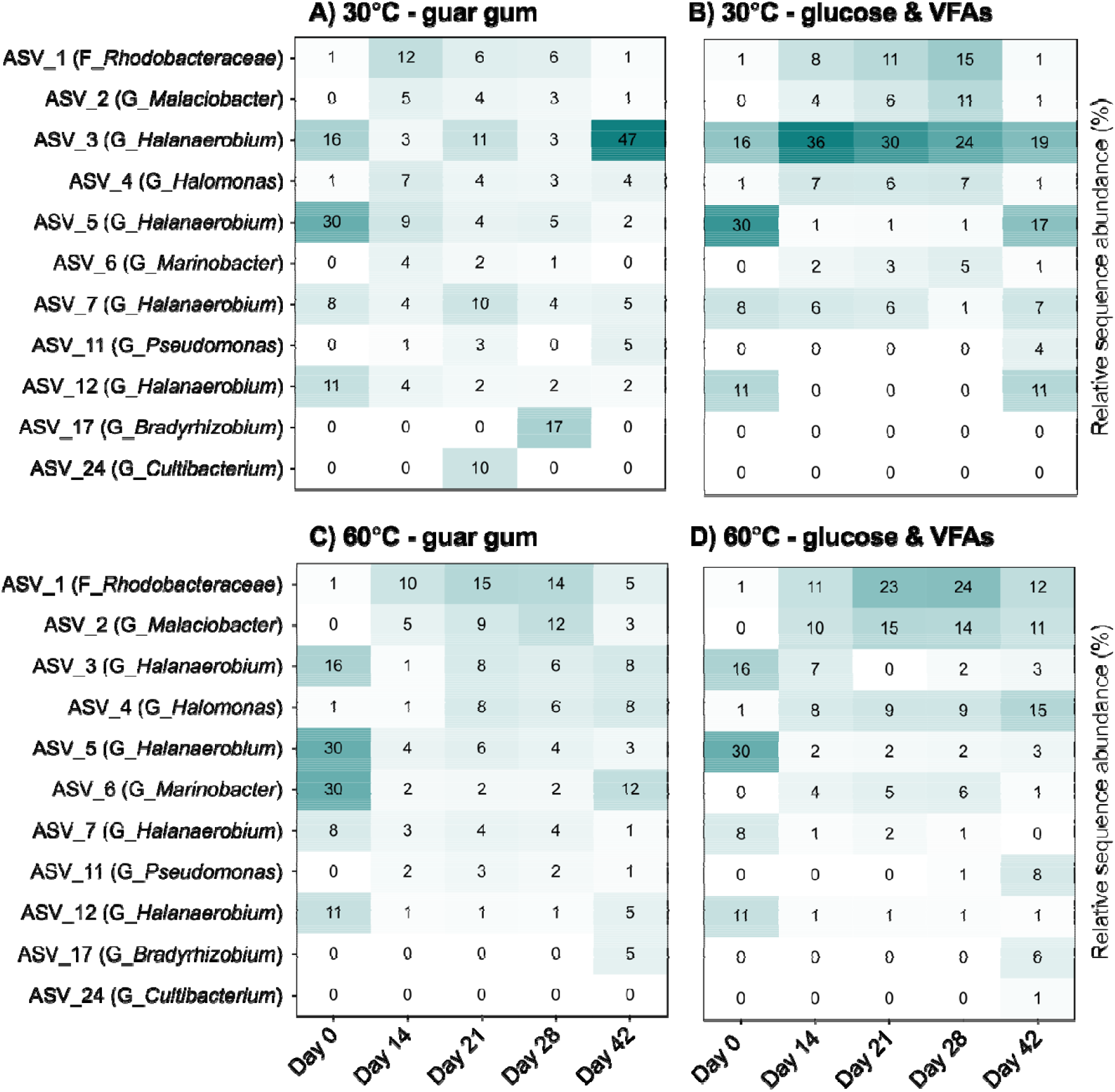
Relative sequence abundance of abundant ASVs during produced water incubations at temperatures mimicking topsides storage ponds (30°C) and subsurface oil reservoirs (60°C). Only ASVs detected at least once at over 5% relative abundance are included. Taxonomy of each ASV is denoted in parentheses based on genus (G) or family (F) level affiliations. Values and shading indicate the average percentage of relative abundance of each ASV at a given time point, based on amplicon sequencing of triplicate incubations.

Putative sulfate-reducing bacteria were observed to increase from low levels at day 0 only in the 60°C enrichments amended with glucose and VFA (Figure 3), corresponding with sulfate reduction to sulfide (Figure 1D). In this enrichment, increases in *Desulfovibrionales* and *Desulfobulbales* orders were observed after 14 days of incubation, reaching 0.9% and 1.9%, respectively (Figure 3A). Within the *Desulfovibrionales*, the most prevalent genus was *Desulfohalobium* (Figure 3B), and within the *Desulfobulbales* the genera most enriched were *Desulfovibrio* and *Desulfocapsa* (Figure 3C). Other incubation types showed very low levels of putative sulfate-reducing microorganisms (below 0.5% relative abundance, Figure S1).

**Figure 3:**
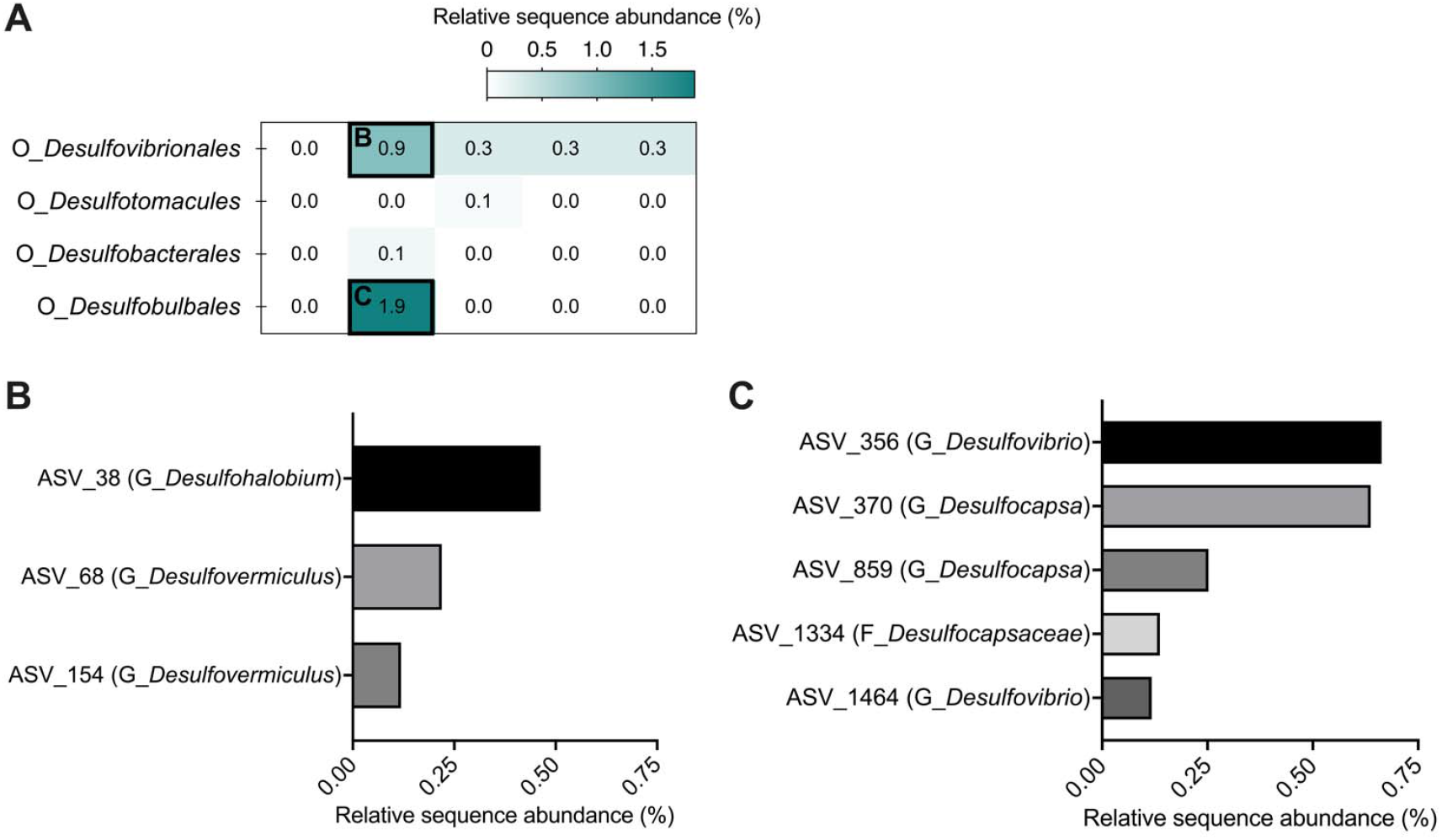
Relative sequence abundance of putative sulfate-reducing bacteria in produced water incubated at 60°C with glucose and volatile fatty acids over a 42-day period. The heatmap (A) shows microbial orders within the samples as the cumulative sequence abundance of ASVs affiliated with these orders throughout the incubation. Histograms show relative sequence abundance of individual ASVs at finer taxonomic resolution within the orders *Desulfovibrionales* (B) and *Desulfobulbales* (C) corresponding to 14 days of incubation when an increase in sulfide was observed (Figure 1D).

### Metagenomic analysis and quality assessment of reconstructed of MAGs

Metagenomic sequencing was performed on DNA extracted from the sample collected at Day 28 from the 30°C guar-amended incubations and from the 60°C incubations amended with glucose and VFA. A total of 25 unique medium- and high-quality metagenome-assembled genomes (MAGs) were reconstructed: 9 from 30°C enrichments and 16 from 60°C enrichments (Table 1, Figure S2, Figure S3). MAGs belonging to the family *Halanaerobiaceae* were only retrieved from the 30°C guar gum enrichments (MAGs 4, 6 and 8; Table 1, Figure 4). Among these, MAG 6 was classified to the genus level (*Halanaerobium*) and MAGs 4 and 8 were classified to the species level as *H. saccharolyticum* and *H. congolense*, respectively. MAGs taxonomically affiliated with lineages capable of sulfate reduction were only found within the enrichments at 60°C; MAGs 10 and 11 both belong to the order *Desulfovibrionales* and could be further classified to the order *Desulfovibrionaceae* (MAG 11) and to the species level for *Desulfohalobium retbaense* (MAG 10).

**Table 1:** Taxonomy and predicted replication rate for 25 metagenome-assembled genomes.

| MAG | Temp. | Taxonomy |  |  |  |  |  |  | Predicted replication rate |
| --- | --- | --- | --- | --- | --- | --- | --- | --- | --- |
|  |  | Domain | Phylum | Class | Order | Family | Genus | Species |  |
| 1 | 30°C | Bacteria | Patescibacteria | Gracilibacteria | BD1-5 | UBA6164 | UBA6164 | - | NA |
| 2 |  | Bacteria | Proteobacteria | Gammaproteobacteria | Pseudomonadales | Pseudomonadaceae | Pseudomonas | *P. stutzeri* | NA |
| 3 |  | Bacteria | Proteobacteria | Alphaproteobacteria | Rhodobacterales | Rhodobacteraceae | Roseovarius | - | NA |
| 4 |  | Bacteria | Firmicutes | Halanaerobiia | Halanaerobiales | Halanaerobiaceae | Halanaerobium | *H. saccharolyticum* | NA |
| 5 |  | Bacteria | Bacteroidota | Bacteroidia | Bacteroidales | Marinilaniaceae | - | - | NA |
| 6 |  | Bacteria | Firmicutes | Halanaerobiia | Halanaerobiales | Halanaerobiaceae | Halanaerobium | - | NA |
| 7 |  | Bacteria | Proteobacteria | Gammaproteobacteria | Pseudomonadales | Halomonadaceae | Halomonas | *H. ventosae* | NA |
| 8 |  | Bacteria | Firmicutes | Halanaerobiia | Halanaerobiales | Halanaerobiaceae | Halanaerobium | *H. congolense* | 2.05 |
| 9 |  | Bacteria | Proteobacteria | Alphaproteobacteria | Rhodobacterales | Rhodobacteraceae | Roseovarius | - | NA |
| 10 | 60°C | Bacteria | Desulfobacterota | Desulfovibrionia | Desulfovibrionales | Desulfohalobiaceae | Desulfohalobium | *D. retbaense* | 2.56 |
| 11 |  | Bacteria | Desulfobacterota | Desulfovibrionia | Desulfovibrionales | Desulfovibrionaceae | - | - | NA |
| 12 |  | Bacteria | Bacteroidota | Bacteroidia | Flavobacterales | Flavobacteraceae | Psychroflexus | *P. salarius* | NA |
| 13 |  | Bacteria | Proteobacteria | Alphaproteobacteria | Rhizobiales | Xanthobacteraceae | Bradyrhizobium | sp003020075 | 1.32 |
| 14 |  | Bacteria | Proteobacteria | Gammaproteobacteria | Pseudomonadales | Halomonadaceae | Halomonas | *H. ventosae* | NA |
| 15 |  | Bacteria | Bacteroidota | Bacteroidia | Bacteroidales | Proxilibacteraceae | SKHV01 | - | NA |
| 16 |  | Bacteria | Bacteroidota | Bacteroidia | Bacteroidales | UBA12170 | - | - | NA |
| 17 |  | Bacteria | Campylobacterota | Campylobacteriia | Campylobacteriales | Arcobacteraceae | - | - | NA |
| 18 |  | Archaea | Halobacteriota | Methanosarcina | Methanosarcinales | Methanosarcinaceae | Methanohalophilus | - | NA |
| 19 |  | Bacteria | Thermotogota | Thermotogae | Petrotogales | - | - | - | NA |
| 20 |  | Bacteria | Patescibacteria | Gracilibacteria | Abconditabacterales | X112 | - | - | NA |
| 21 |  | Bacteria | Defferibacterota | Defferibacteres | Defferibacteriales | Flexistipitaceae | Flexistipes | - | NA |
| 22 |  | Bacteria | Proteobacteria | Alphaproteobacteria | Rhodobacterales | Rhodobacteraceae | Roseovarius | - | 1.62 |
| 23 |  | Bacteria | Firmicutes | Bacilli | Izemoplasmales | Izemoplasmataceae | - | - | NA |
| 24 |  | Bacteria | Bacteroidota | Bacteroidia | Flavobacterales | Flavobacteraceae | Psychroflexus | - | NA |
| 25 |  | Bacteria | Patescibacteria | Gracilibacteria | BD1-5 | UBA6164 | UBA6164 | - | 2.50 |

**Figure 4:**
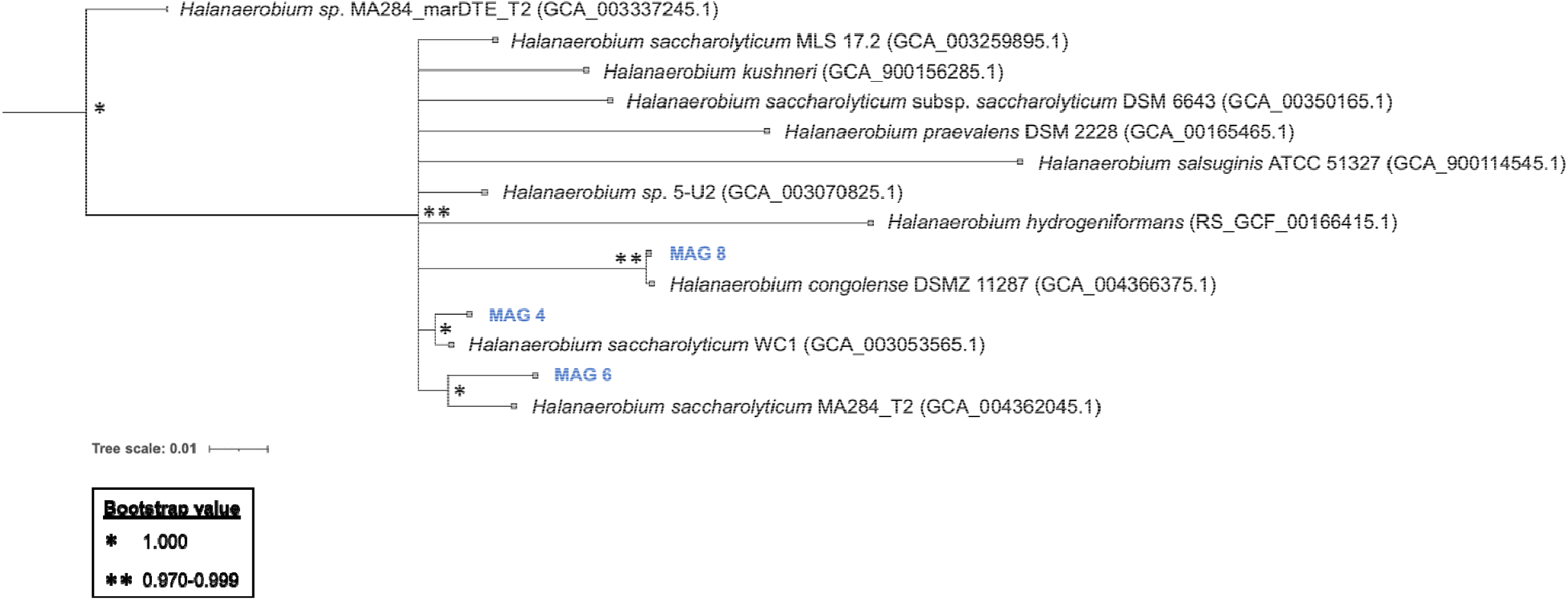
Phylogenomic relationships among *Halanaerobiaceae* from Permian Basin produced water and other environments reconstructed from GTDK-Tk. Three *Halanaerobium* MAGs from this study (i.e., 30°C incubation with guar gum) are highlighted in bold blue font. GenBank accession numbers for genomes from other environments are indicated in parentheses. These include the type strain *Halanaerobium congolense* (22) and different *H. saccharolyticum* strains that are closely related to new MAGs uncovered in this study. Tree scale is shown at the bottom left of the figure.

Sulfate-reducing *D. retbaense* (MAG 10) had the highest replication rate among five MAGs, which also included *Halanaerobium congolense* MAG 8, as indicated in Table 1.

## Metabolic features encoded in MAGs

### Guar gum fermentation potential in MAGs from 30°C incubations

Guar gum biodegradation by extracellular enzymes involves the ß-1,4-mannanase catalyzing hydrolysis of the mannose backbone of the polymer and the α -1,6-galactosidase breaking down galactose units branching out of the mannose backbone (14). Both enzymes are encoded by *Halanaerobium* MAG 6, with *Halanaerobium* MAGs 4 and 8 also encoding the alpha-galactosidase. The beta-mannanase was also found in *Marinilabillaceae* MAG 5. ABC transporters for internalization of mannose are encoded by all three *Halanaerobium* MAGs as well as by *Pseudomonas* MAG 2. The mannose PTS system to internalize and phosphorolyze mannose to mannose-6-phosphate was found in *Halanaerobium* MAG 8. Galactose ABC transporter genes were found in all three *Halanaerobium* MAGs 4, 6 and 8, as well as in *Halomonas* MAG 7.

The mannose PTS system (EIIA component) is known to internalize and phosphorolyze mannose to mannose-6-phosphate (found in MAG 6), which can then be further converted to fructose-6-phosphate by the mannose-6-phosphate isomerase and introduced in the glycolysis pathway. All three *Halanaerobium* genomes encode mannose-6-phosphate isomerases and enzymes needed for glycolysis (Figure 5). Mannose conversion to mannose-6-phosphate requires a hexokinase that is not in any of the MAGs retrieved from the 30°C guar gum enrichments. *Halanaerobium* MAGs 6 and 8 have complete sets of genes for the Leloir pathway to ferment galactose to UDP-glucose. Further conversion of UDP glucose to glucose-1-phosphate by the UTP-glucose-1-phosphate-uridyltransferase, and glucose-6-phosphate by phosphoglucomutase (to integrate into glycolysis) are also encoded in both genomes (Figure 5).

**Figure 5:**
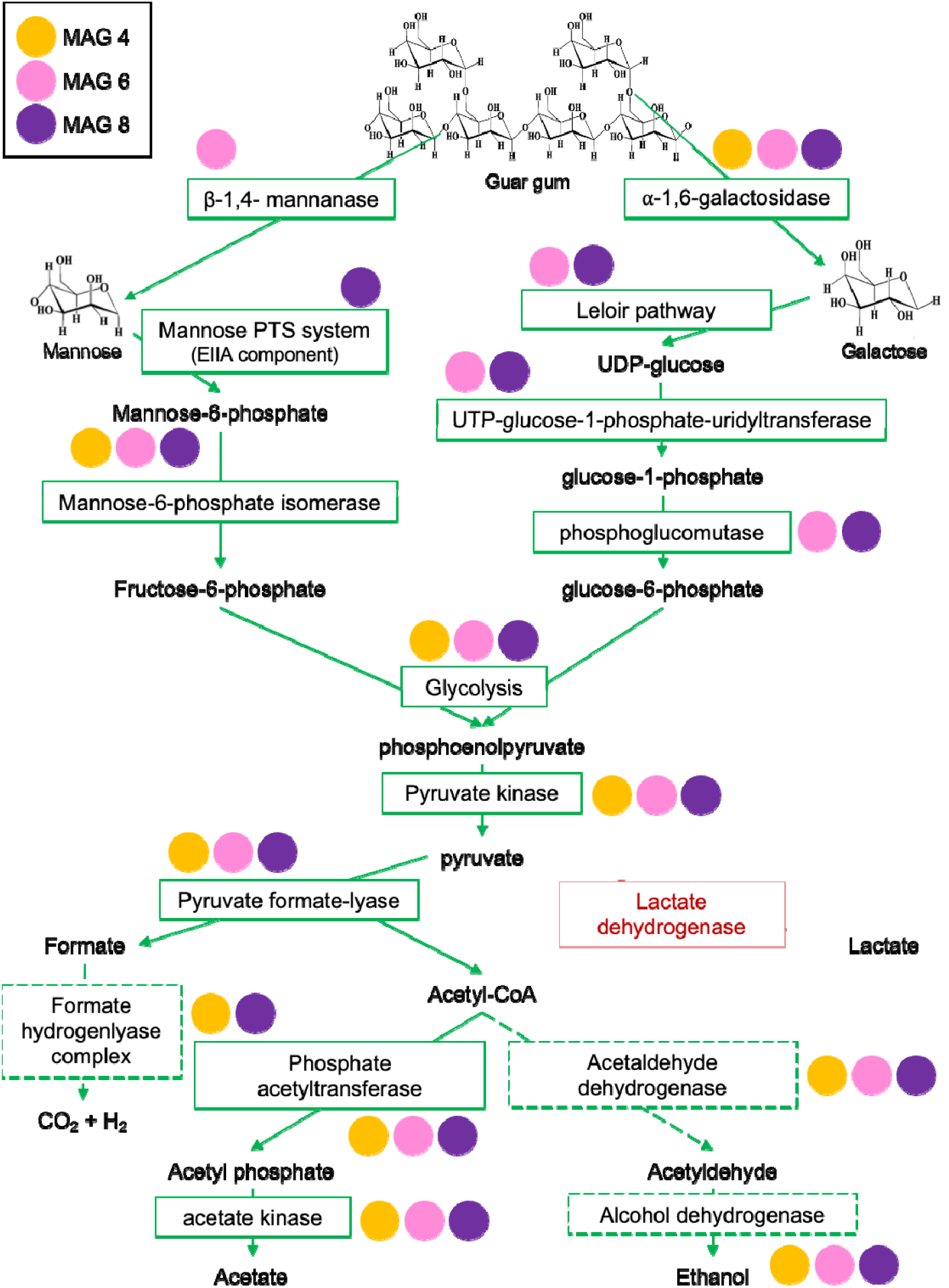
Genes present within three *Halanaerobium* genomes (Figure 4) encoding enzymes for catalyzing biodegradation of guar gum polymers to acetate and CO_2_ through the mixed-acidfermentation pathway. Green arrows and boxes indicate genes and pathways detected in *Halanaerobium* MAGs 4, 6 and 8 that were enriched at 30°C in the presence of guar gum. Red arrows and boxes signify non detection. Dashed green lines indicate partial detection of genes but not the complete pathway.

### Mixed-acid fermentation potential in MAGs from 30°C incubations

Mixed-acid fermentation includes production of lactate, acetate, ethanol, CO_2_ and H_2_ through the oxidation of pyruvate. Genes for lactate dehydrogenase were not detected in three *Halanaerobium* MAGs or in any contigs annotated from the assemblies prior to binning, whereas genes for acetate production were found in all three *Halanaerobium* MAGs (Figure 5). Furthermore, although complete pathways were not observed, genes for the production of CO_2_ and H_2_ are present in *Halanaerobium* MAGs 4 and 8. Genes encoding enzymes for ethanol production were observed in all three *Halanaerobium* MAGs.

### Potential for sulfur metabolism in MAGs from 30°C and 60°C incubations

Members of the genus *Halanaerobium* have been studied for their potential to catalyze sulfidogenesis via thiosulfate reduction, yet no genes related to thiosulfate metabolism were found in *Halanaerobium* genomes recovered from 30°C incubations with guar gum (or any assembled contigs). This includes genes involved in the SOX system to reduce thiosulfate to sulfate, thiosulfate reductase (*phsABC*), tetrathionate reductase, thiosulfate sulfurtransferase and rhodanese genes; despite these genes being present in other *Halanaerobium* spp., they were not detected in MAGs 4, 6 and 8, which all exhibited over 89% estimated completeness (Table 1, Figure S3). Genes encoding the complete pathway for dissimilatory sulfate reduction were identified in genomes assembled from the 60°C incubations classified as *Desulfohalobium* (MAG 10) and *Desulfovibrionaceae* (MAG 11). Genes for dissimilatory sulfate reduction were not detected in the metagenomes associated with incubation at 30°C.

### Organotrophic potential of thermophilic sulfate-reducing bacteria at 60°C

Genes encoding complete pathways for the oxidation of acetate to CO_2_ and acetyl-coA (acetyl-coA pathway) in MAGs 10 and 11 with higher replication rates (Table 1) suggest that these organisms can couple acetate oxidation to sulfate reduction at higher temperatures (60°C) in the subsurface. Highly incomplete genomic pathways for the TCA cycle were retrieved from both MAG 10 and 11 with key steps missing (genes for the conversion of oxaloacetate to citrate, succinate to fumarate, fumarate to malate and malate to oxaloacetate). Interestingly, while no genes related to hydrolysis of guar gum were found within these genomes, genes for the internalization of mannose and conversion to mannose-6-phosphate (PTS system EIIA components) were retrieved from sulfate-reducing *Desulfohalobium* MAG 10 and *Desulfovibrionaceae* MAG 11, with the latter also having hexokinase genes for converting mannose to mannose-6-phosphate. Both sulfate-reducing microorganisms have genes for the conversion of mannose-6-phosphate to fructose-6-phosphate, and genes for the glycolysis pathway that are complete (MAG 10) or nearly complete (MAG 11). This indicates the potential to use mannose to complete glycolysis, coupled to sulfate reduction.

## Discussion

### Fermentative metabolism under lower temperature storage pond conditions

Members of the genus *Halanaerobium* have been extensively documented within hydraulically fractured shale oil reservoirs where they potentially catalyze fermentation (notably guar gum utilization) and thiosulfate reduction (4–7, 26). Despite *Halanaerobium* being routinely recovered from shale formations with downhole temperatures up to 60°C (10), *Halanaerobium* species have not been cultivated above 42°C (5, 26). Consistent with these observations, *Halanaerobium* ASVs and metagenomic raw reads increased in relative abundance during incubations at 30°C with guar gum but not in 60°C incubations (Figure 2, Table 1, Figure 5 and Figure S2). This is corroborated by *Halanaerobium* MAGs 4, 6 and 8 that were enriched at 30°C all having optimal growth temperatures of 39-40°C estimated by the Tome algorithm (27).

Previous studies have not reported on biochemical pathways for guar gum metabolism by *Halanaerobium* spp., although isolated members of this genus have been shown to biodegrade the polymer, commonly used as a gelling agent during hydraulic fracturing operations (5, 12-15). Liang et al. (2016) isolated a strain of *Halanaerobium* able to couple guar gum conversion to acetate with the reduction of thiosulfate to sulfide (5). In the present study, *Halanaerobium* MAGs encode mannanase and galactosidase genes for internalizing mannose and galactose derived from extracellular hydrolysis of guar gum, i.e., ABC transporters for both sugars or the mannose PTS system (Figure 5). These bacteria can then metabolize these sugars through a mixed-acid fermentation pathway (Figure 5) to produce acetate and CO_2_, which were both observed to accumulate in the corresponding enrichment cultures (Figure 1A).

Accordingly, fermentation of larger substrates by *Halanaerobium* (Figure 5) may be a more plausible metabolism than thiosulfate reduction in lower-temperature topsides conditions like produced water storage ponds. Fermentation appeared to be the predominant metabolism when produced water was incubated under temperatures representative of storage ponds (Figure 1), and the communities were dominated by *Halanaerobium* (Figure 2). The absence of thiosulfate reduction genes in three near-complete *Halanaerobium* genomes is notable. In particular, *Halanaerobium* MAG 8 shares 97.8 average nucleotide identity with type strain *H. congolense* that has been characterized as a thiosulfate reducer (22). To our knowledge, only one study has measured the presence of thiosulfate in produced water where *Halanaerobium* with genes for thiosulfate metabolism was also detected, however the thiosulfate concentration of 0.02 mM was very low (6). *Halanaerobium* strain DL-01 referred to above for its ability to couple thiosulfate reduction with guar metabolism was shown to do so in the presence of 10 mM thiosulfate amended to Barnett shale produced water in which thiosulfate levels were otherwise below detection limits. Other studies of shale reservoir *Halanaerobium* found genes for the reduction of thiosulfate but did not report thiosulfate concentrations in the produced water (7, 21, 28). Based on these observations combined with results reported here, *in situ* thiosulfate reduction by *Halanaerobium* is difficult to confirm. More work is needed to detect and measure levels of thiosulfate in these systems to understand its potential role as a precursor for sulfide biogenesis thereby implicating *Halanaerobium* directly in souring and associated corrosion problems.

### *Halanaerobium* acting in concert with SRB during produced water re-use

Sulfate reduction to sulfide at 60°C in incubations amended with glucose and volatile fatty acids (Figure 1) appears to be catalyzed by thermophilic sulfate-reducing bacteria including *Desulfohalobium* MAG 10 and *Desulfovibrionaceae* MAG 11, despite amplicon sequencing suggesting these groups were present in relatively low abundance (Figure 3) during the incubation period. Sulfate reduction corresponding to low abundances of putative sulfate-reducing microorganisms have been reported in other environmental systems (29, 30). In the experiments presented here, sulfate reduction took place during the first half of the 42-day incubation period (Figure 1D), coinciding with the largest increase in relative sequence abundance of ASVs affiliated with known sulfate-reducing bacteria (Figure 3). Reconstruction of *D. retbaense* (MAG 10) and a member of the *Desulfovibrionaceae* family (MAG 11) showed that in addition to complete pathways for dissimilatory sulfate reduction, these genomes contained incomplete sets of genes for the TCA cycle and all genes required for acetate oxidation via the acetyl-coA pathway.

This understanding is consistent with *Halanaerobium* spp. being detected in saline oil fields experiencing high levels of souring caused by sulfate-reducing microorganisms (32, 33). Metagenomic sequencing points to the main role of *Halanaerobium* in this system being able to secrete extracellular enzymes to hydrolyze guar gum, thereby releasing mannose and galactose and leading to the release of other by-products (hydrogen, carbon dioxide and acetate) for other microorganisms to utilize whether in topsides storage (e.g., storage ponds) mimicked by 30°C enrichments or in the subsurface mimicked by 60°C enrichments. This evidence of fermentative metabolism is particularly important in the context of re-injecting produced water after a period of topside storage at lower temperature that affords *Halanaerobium* the opportunity to generate electron donors for sulfate reduction, including hexose sugars (mannose, galactose), acetate, ethanol and hydrogen. If these compounds are introduced into the subsurface in the presence of sulfate, thermophilic sulfate-reducing microorganisms in the reservoir are presented with the opportunity to catalyze reservoir souring via dissimilatory sulfate reduction. Supporting this statement is the presence of acetate and sulfate within the sample obtained from the Permian Basin oil reservoir produced water obtained for this study (Table S1). Also, extracellular ß-1,4-mannanase are secreted within the produced water under storage pond temperatures to hydrolyze guar gum leading to increased levels of mannose (*Halanaerobium* MAGs 4, 6 and 8), which could be directly coupled to dissimilatory sulfate reduction by *Desulfohalobium* (MAG 10) and a *Desulfovibrionaceae* (MAG 11).

In conclusion, produced water incubations at 30°C and 60°C highlight the potential for cooperative metabolic activity by *Halanaerobium* and sulfate-reducing microorganisms via produced water storage and reuse in hydraulically fractured shale formations. The importance of organotrophic metabolism by *Halanaerobium* and its potential for promoting reservoir souring by providing substrates for sulfate-reducing microorganisms was illustrated. Fermentative metabolism is the most likely explanation for the prevalence of *Halanaerobium* in shale oil fields.

## Experimental procedures

### Microbial enrichments

Produced water samples from Permian Basin hydraulically fractured wells (New Mexico, USA) were used to set up microbial enrichments. Enrichment medium (per 1L) was composed of: 1g NH_4_Cl, 10g MgCl_2_•6H_2_O, 0.1g CaCl_2_•2H_2_O, 1g KCl, 100g NaCl, 0.72g Cysteine HCl, 500 μM K_2_HPO_4_/KH_2_PO_4_, 0.2% (v/v) NAHCO_3_, 10 mL trace element solution and 10 mL vitamin solution (6). Microcosms were prepared by combining 20 mL produced water with 80 mL of medium in 125-ml serum bottles under anoxic conditions. Serum bottles were immediately sealed with sterile rubber stoppers and the headspace was exchanged with N_2_:CO_2_ (90:10%). Two different substrate types were used, either glucose (10 mM) together with a volatile fatty acid (VFA) mixture (acetate, butyrate, formate, lactate, propionate, succinate; each at 5 mM) or guar gum at a concentration of 0.05 % (w/v). Sulfate was added to all microcosms to a final concentration of 10 mM. Bottles were incubated at 30ºC or 60ºC. Sterile controls (medium only with substrates) and substrate-free controls (medium and produced water without substrates) were prepared and incubated in parallel. Subsamples from microcosms were removed using N_2_:CO_2_-flushed sterile syringes and immediately stored at -20°C until further analysis.

### Concentrations of metabolic reactants and products

Sulfate depletion was assessed by ion chromatography (Dionex ICS-5000) using an analytical column (AS23) with 8 mM Na_2_CO_3_/ 1 mM NaHCO_3_ eluent at a flow rate of a 1 mL/min. Chromeleon software was used for visualization. Peak area calibration used Na_2_SO_4_ standard solutions. Sulfide production was measured using a spectrophotometric assay (34). Samples for sulfide analysis were collected at different timepoints of the experiment and mixed with zinc acetate (20% (v/v)) to limit sulfide volatility (35). Samples were kept frozen until analyzed.

Volatile fatty acid levels were measure using high-performance liquid chromatography (HPLC) in an Ultimate 3000 RSLC system with a 5 mM H_2_SO_4_ mobile phase at a flow rate of 0.6 mL/min and a temperature of 60ºC using an Bio-Rad Aminex HPX-87H column. Carbon dioxide production was measured using gas chromatography with a flame ionization detector (GC-FID) by injecting headspace gas into an Agilent 7890B gas chromatoghraph equipped with a Hayesep N packing column (stainless steel tubing, 0.5 meter length × 1/8 inch outer diameter × 2 mm internal diameter, mesh size 80/100). Helium was used as a carrier gas (flow rate (21 mL/min). Oven temperature was set to 105°C and carbon dioxide was detected by a thermal conductivity detector set to 200°C.

### Genomic DNA extraction

5 mL aliquots were removed from microcosms at day 0, 7, 14, 21 and 28 and then frozen until genomic DNA was extracted using the DNeasy PowerSoil extraction kit (Qiagen) following the manufacturer’s instructions. Resulting DNA concentrations were measured by fluorimetry (Qubit, Qiagen). DNA was used for both 16S rRNA gene amplicon sequencing and shotgun metagenomic sequencing, as described below.

### 16S rRNA gene amplicon sequencing and sequence analysis

The V4 hypervariable region of the 16S rRNA gene was amplified using primers 515F (GTGYCAGCMGCCGCGGTAA) and 806R (GGACTACNVGGGTWTCTAAT). PCR reactions with a total volume of 25 μL included: 2× KAPA HiFi Hot Start Ready Mix (KAPA Biosystems), a final concentration of 0.1 mM for each primer and 7.5 μL of DNA sample. PCR included an initial denaturation at 95ºC for 5 minutes followed by 30 cycles of denaturation at 95ºC for 30 seconds, annealing at 55ºC for 45 seconds and extension at 72ºC for 1 min. This was followed by a final extension for 5 minutes at 72ºC. PCR products followed post-PCR Triplicate PCR products were pooled prior to cleanup and indexing. Indexed amplicon samples were sequenced using Illumina’s v3 600-cycle (paired-end) reagent kit on an in-house Illumina MiSeq benchtop sequencer (36) after all DNA extraction blanks and PCR reagent blanks were confirmed negative for amplification. Quality-controlled reads were merged and subsequently dereplicated to construct an amplicon sequence variant (ASV) table. Relative abundance calculations were based on the number of unrarefied reads per sample. Analysis and visualization were done using R and GraphPad Prism (37, 38).

### Metagenomic sequencing

Genomic DNA extracted from produced water enrichments after 28 days of incubation was sent to the Centre for Health Genomics and Informatics (University of Calgary, Calgary, Canada) for library preparation using sonication (Covaris) and sequencing using a NovaSeq platform (Illumina). Adapters were removed, the last 151 bp of the last read were trimmed, contaminants (PhiX control V3 library, Illumina) were filtered and low quality ends were clipped off with BBDuk (39). Assembly was done with Megahit using a co-assembly for replicates of a given sample, and reads were assembled using Bowtie2 (40, 41). DasTool allowed for binning of the contigs while combining outputs from Concoct, Maxbin and Metabat (42–45). Quality assessment of the bins was performed with CheckM ‘lineage_wf’ (v.1.1.2) with taxonomy assigned using GTDB-Tk ‘classify_wf’ (v.1.5.0) (46, 47). Only bins with <10% and completeness >70% were retained for annotation and further study. DRAM (v.1.2) was used to annotate MAGs for functional potential (Colorado State University, CO, United States; Shaffer et al., 2020). PhyloFlash allowed for taxonomic identification from the metagenomic reads (49). Estimations of temperature optima for metagenome-assembled genomes (MAGs) used Tome (27). iRep allowed for estimation of replication rates on MAGs (50). Genomes outside the minimal requirements (under 2% contamination, over 75% complete) to analyze with iRep were excluded from the analyses.

### Phylogeny

Phylogenomic trees were constructed using CheckM version 1.1.3 and included 2,052 finished and 3,604 draft genomes from the Integrated Microbial Genomes (IMG) database (46). Genes were aligned using GAMMA and WAG models. Nodes were interpreted as bootstrap values and the tree was constructed using Dendroscope for visualization (51).

## Conflict of Interest

The authors declare no conflict of interest.

## Author Contributions

P.E. and C.R.J.H. secured produced water samples, G.S. and A.C. designed the study, G.S. performed lab work and G.S. wrote the manuscript with input from C.R.J.H. All authors have read and agreed to the published version of the manuscript.

## Funding

This work was supported by grants the Natural Sciences and Engineering Research Council of Canada (NSERC) to C.R.J.H., and by scholarship funding to G.S. from NSERC (Alexander Graham Bell scholarship program) and the Eyes High program (University of Calgary).

## Acknowledgments

The authors would like to acknowledge that this work was done on the traditional territories of the Blackfoot Confederacy (Siksika, Kainai, Piikani), the Tsuut’ina, the Îyâxe Nakoda Nations, the Métis Nation (Region 3), and all people who make their homes in the Treaty 7 region of Mohkínsstsisi (now named Calgary). The authors would like to thank Dana Jwad and Rohan Khan for laboratory support as well as Srijak Bhatnagar for bioinformatics support. This work was funded by grants from Chevron and the Natural Sciences and Engineering Research Council of Canada to CRJH, and by the Eyes high doctoral scholarship (University of Calgary) and the Alexander Graham Bell scholarship (NSERC) to GS.

## Data Availability Statement

The data discussed in this manuscript is available online under the Genbank bioproject number PRJNA972118.

## Notes

### Competing Interest Statement

The authors have declared no competing interest.

