## Supplementary_figures-Tables for "Polymer biodegradation by *Halanaerobium* promotes reservoir souring during hydraulic fracturing"

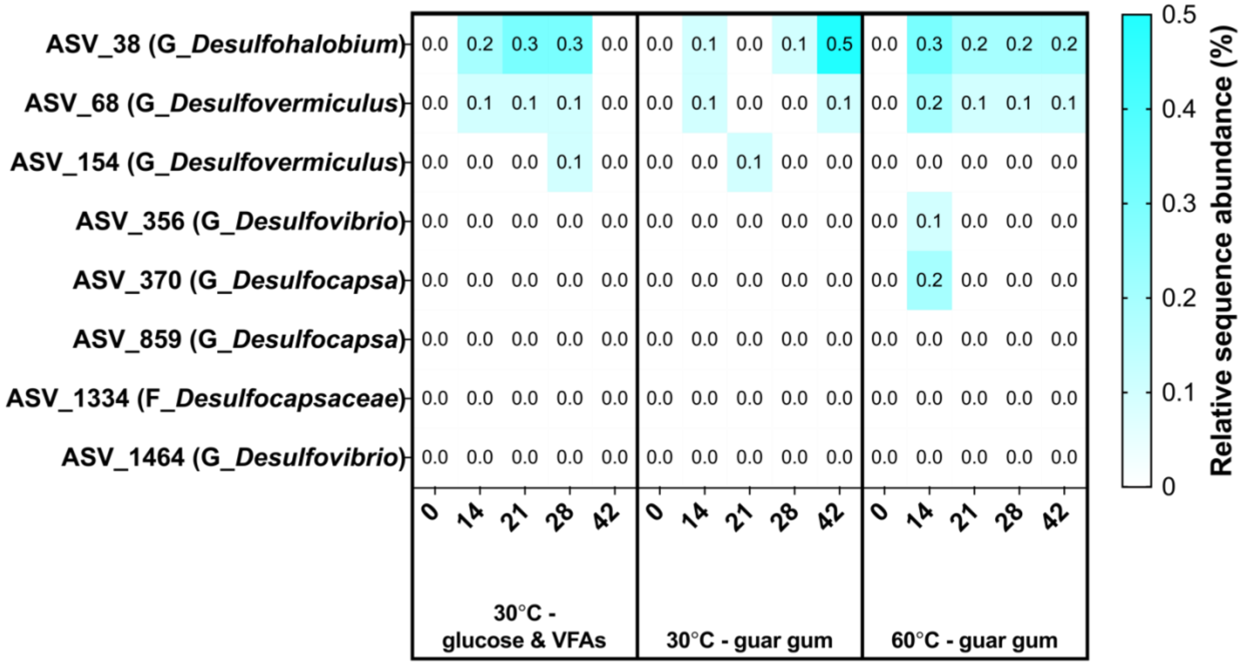


**Figure S1:** Relative sequence abundance of putative sulfate-reducing bacteria in produced water incubated at 30°C or 60°C with guar gum or glucose and volatile fatty acids over a 42-day period. The heatmap shows the cumulative relative abundance of ASVs throughout the incubation.

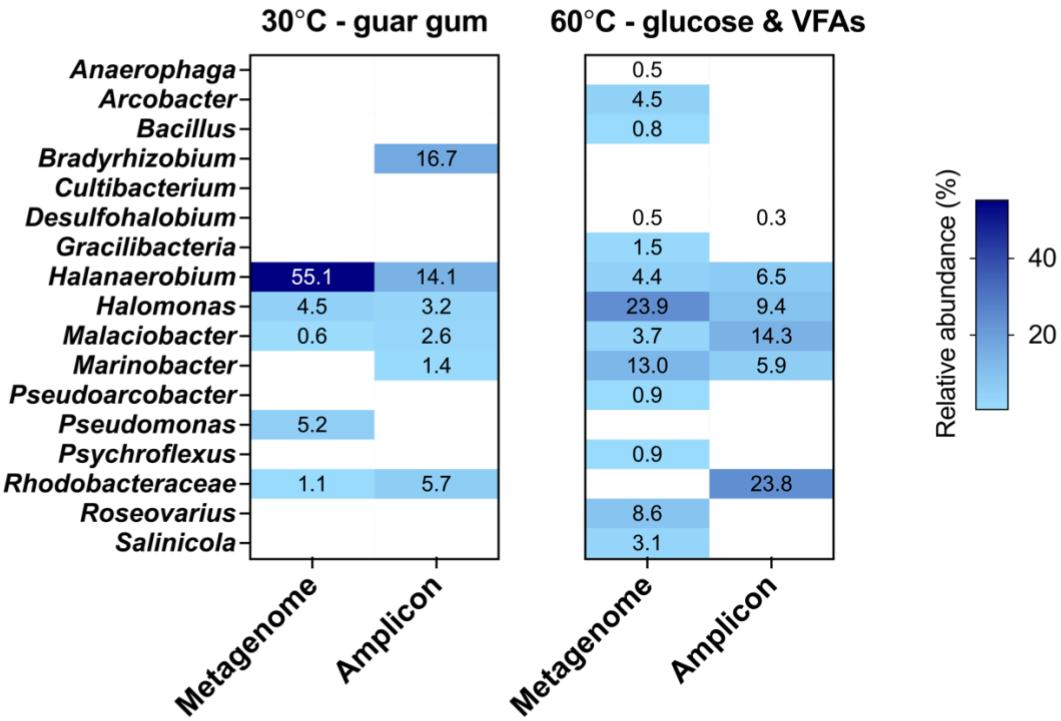


**Figure S2:** Microbial community composition based on metagenome reads at 28 days (labeled "metagenome") of incubation from samples mimicking topsides temperature (30°C; supplemented with guar gum) and subsurface oil reservoirs temperature (60°C; supplemented with glucose and VFA) compared to amplicon sequencing data (labeled "amplicon", Figure 2). Only ASVs over 0.5% relative abundance in either group of enrichments are included.

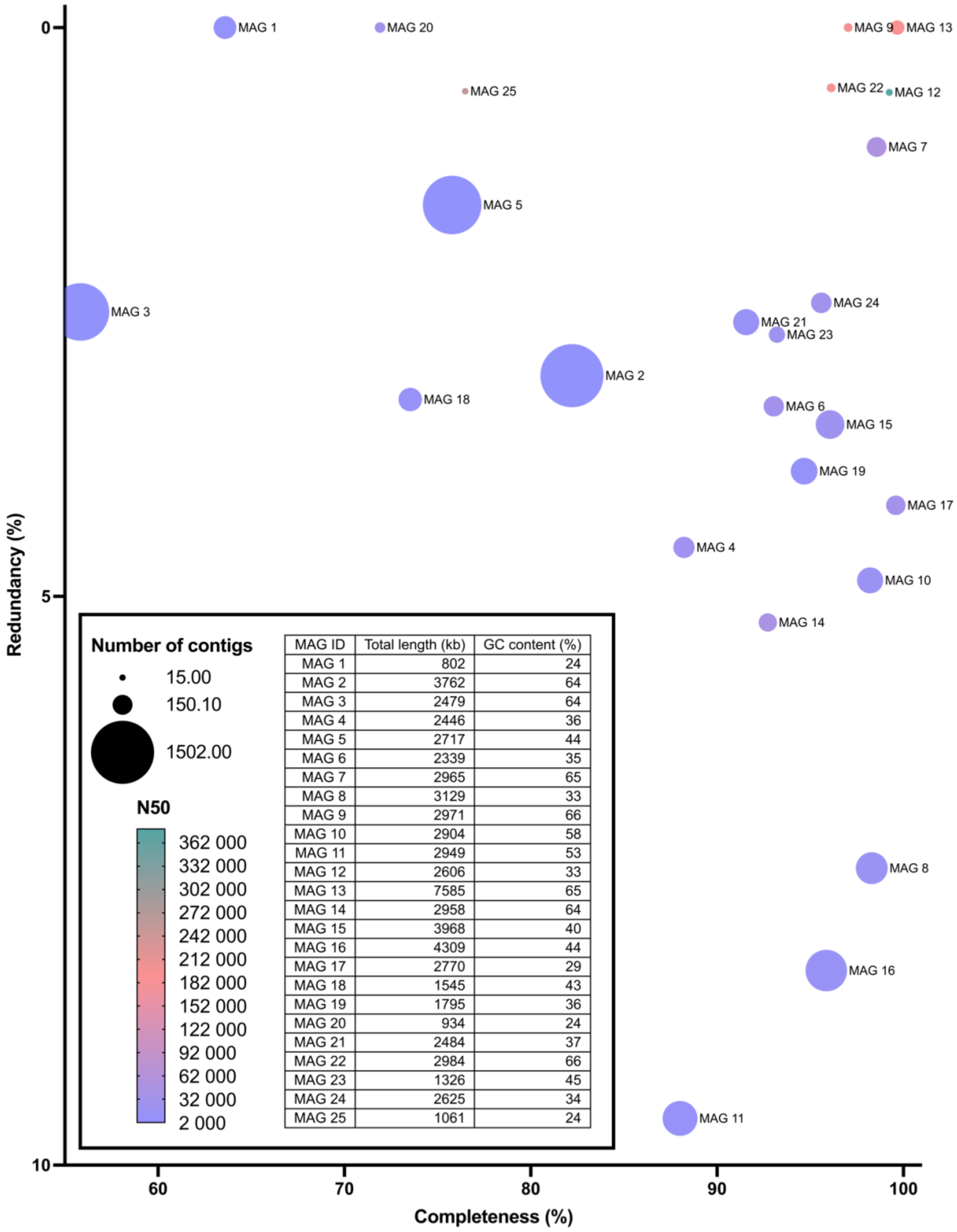


| MAG ID | Total length (kb) | GC content (%) |
|---|---|---|
| MAG 1 | 802 | 24 |
| MAG 2 | 3762 | 64 |
| MAG 3 | 2479 | 64 |
| MAG 4 | 2446 | 36 |
| MAG 5 | 2717 | 44 |
| MAG 6 | 2339 | 35 |
| MAG 7 | 2965 | 65 |
| MAG 8 | 3129 | 33 |
| MAG 9 | 2971 | 66 |
| MAG 10 | 2904 | 58 |
| MAG 11 | 2949 | 53 |
| MAG 12 | 2606 | 33 |
| MAG 13 | 7585 | 65 |
| MAG 14 | 2958 | 64 |
| MAG 15 | 3968 | 40 |
| MAG 16 | 4309 | 44 |
| MAG 17 | 2770 | 29 |
| MAG 18 | 1545 | 43 |
| MAG 19 | 1795 | 36 |
| MAG 20 | 934 | 24 |
| MAG 21 | 2484 | 37 |
| MAG 22 | 2984 | 66 |
| MAG 23 | 1326 | 45 |
| MAG 24 | 2625 | 34 |
| MAG 25 | 1061 | 24 |

**Figure S3:** Quality of 25 bins retrieved from the Permian Basin produced water enrichments after 42 days of incubations.

**Table S1:** Concentration of various compounds measured within the Permian Basin produced water used for microbial enrichments in this study.

| **Compound** | **Concentration (mM)** |
|---|---|
| Sulfate | 4.1 |
| Thiosulfate | Below detection limits |
| Acetate | 2.1 |
| Butyrate | Below detection limits |
| Formate | ND |
| Lactate | Below detection limits |
| Propionate | 0.2 |
| Succinate | ND |

**Table S2:** Optimal growth temperature computationally predicted using Tome for three different Halanaerobium MAGs found within the 30°C enrichments with guar gum.

| **MAG ID** | **Predicted optimal growth temperature (°C)** |
|---|---|
| 4 | 39 |
| 6 | 39 |
| 8 | 40 |
